# A gel-coated air-liquid-interface culture system with tunable substrate stiffness matching healthy and diseased lung tissues

**DOI:** 10.1101/2023.07.29.550710

**Authors:** Zhi-Jian He, Catherine Chu, Riley Dickson, Li-Heng Cai

## Abstract

Since its invention in the late 1980s, the air-liquid-interface (ALI) culture system has been the standard *in vitro* model for studying human airway biology and pulmonary diseases. However, in a conventional ALI system, cells are cultured on a porous plastic membrane that is much stiffer than human airway tissues. Here, we develop a gel-ALI culture system by simply coating the plastic membrane with a thin layer of hydrogel with tunable stiffness matching that of healthy and fibrotic airway tissues. We determine the optimum gel thickness that does not impair the transport of nutrients and biomolecules essential to cell growth. We show that the gel-ALI system allows human bronchial epithelial cells (HBECs) to proliferate and differentiate into a pseudostratified epithelium. Further, we discover that HBECs migrate significantly faster on hydrogel substrates with stiffness matching that of fibrotic lung tissues, highlighting the importance of mechanical cues to human airway remodeling. The developed gel-ALI system provides a facile approach to studying the effects of mechanical cues on human airway biology and in modeling pulmonary diseases.

**New and Noteworthy:** In a conventional air-liquid-interface (ALI) system, cells are cultured on a plastic membrane that is much stiffer than human airway tissues. We develop a gel-ALI system by coating the plastic membrane with a thin layer of hydrogel with tunable stiffness matching that of healthy and fibrotic airway tissues. We discover that human bronchial epithelial cells migrate significantly faster on hydrogel substrates with pathological stiffness, highlighting the importance of mechanical cues to human airway remodeling.

## 1. Introduction

Cell culture models allow for recapitulating essential biological features of human organs or tissues, and thus represent powerful tools for studying cell biology, disease modeling, and drug discovery. For instance, air-liquid interface (ALI) culture system was developed in the late 1980s to study human airway biology and to model pulmonary diseases (1). In an ALI system, human bronchial epithelial cells (HBECs) are cultured on a porous membrane, through which nutrients are transported from cell culture media at the basal side, whereas on the apical side cells are in contact with air (2). The ALI system allows airway basal cells to differentiate into ciliated cells and goblet cells, eventually forming a pseudostratified columnar epithelium that recapitulates essential biological features of the human airway epithelium (3–6) (**Fig. 1a**). Thus, since its invention the ALI culture system represents the standard *in vitro* model for studying human airway defense (7, 8), airway biology (9, 10), airway remodeling (11, 12), airway infection (13, 14), and muco-obstructive lung diseases (15) that include chronic obstructive pulmonary disease (COPD) (16), cystic fibrosis (17, 18), primary ciliary dyskinesia (19), and non–cystic fibrosis bronchiectasis (20).

**Figure 1.**
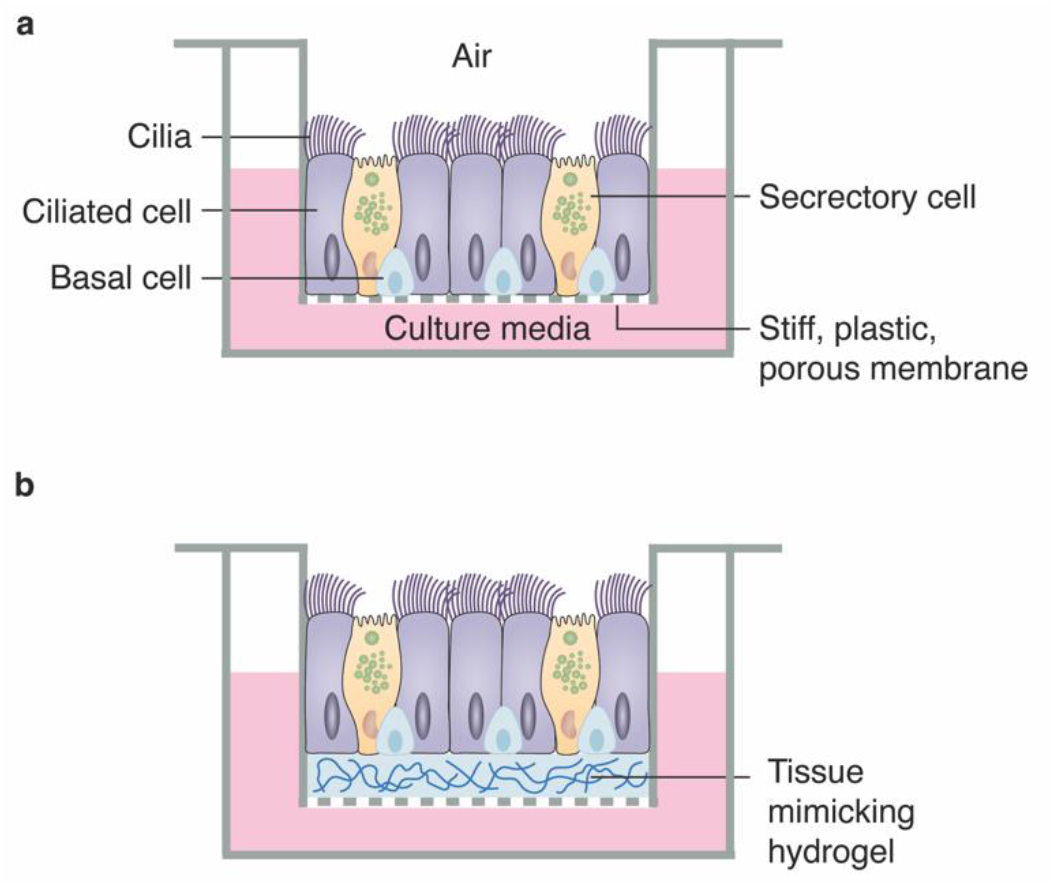
Human bronchial epithelial cell culture systems. **(a)** In a conventional ALI system, HBECs are cultured on a stiff, porous plastic membrane with a fixed stiffness on the order of 10^9^ Pa. **(b)** In a gel-ALI system, HBECs are cultured on a layer of porous hydrogel with tunable stiffness matching that of lung tissues.

The classical ALI culture system, however, suffers from two major limitations. *First*, the porous membrane is made of plastics such as polyester or polyethylene terephthalate, which has a shear modulus on the order of 10^9^ Pa, ∼10^6^ times higher than that of human airway tissues. By contrast, in the human airway, epithelial cells are supported by extracellular matrix (ECM), which not only provides structural support and mechanical stability to the airway epithelium, but also serves as a microenvironment to regulate cell behavior and function (21, 22). For example, substrates with ECM mimicking stiffness affect the morphology of alveolar epithelial cells (23), as well as the size of focal adhesions and the actin cytoskeleton (24). *Second*, the stiffness of the plastic membrane is nearly fixed and cannot be changed. By contrast, the stiffness of the lung tissues varies with pulmonary injuries or diseases (25). In idiopathic pulmonary fibrosis (IPF), a progressive lung disease of unknown causes that is characterized by dysregulated lung remodeling resulting in massive ECM production, the lung tissues are stiffened by nearly ∼16 times from ∼1 kPa in health to 16 kPa in disease (26, 27). Thus, the stiff plastic membrane intrinsically limits the applicability of the classical ALI system in studying the effects of ECM mechanical properties on airway epithelial cell behavior and function.

To better recapitulate the mechanical environment of lung tissues, various materials and methods have been developed to replace or coat the plastic membrane of the classical ALI system. For example, an elastomeric polydimethylsiloxane (PDMS) membrane can be used to separate two microfluidic channels, creating an ALI system for HBEC culture. Because the elastomeric membrane can be reversely deformed under stress, the microfluidic device, or lung-on-a-chip, provides a powerful platform for mimicking the cyclic mechanical strain of the lung alveolar barrier during breathing (28). Moreover, recently the same lung-on-a-chip concept was successfully extended to culture small airway cells (29). However, because PDMS elastomers are not permeable to water and nutrients, microscale pores must be introduced to the elastomeric membrane, making the lung-on-a-chip system dependent on multi-step, sophisticated microfabrication (30). Additionally, the stiffness of typical PDMS elastomers (31) is much higher than that of healthy and fibrotic lung tissues (32). Unlike elastomers, hydrogels are porous networks that allow for good nutrient transport and are very soft, with stiffness tunable in a wide range that can precisely match that of various tissues (33, 34). As a result, the hydrogel is a promising candidate for substituting the plastic membrane in the ALI culture system. For example, an ALI magnetic microboat culture system has recently been developed (35), in which a layer of polyacrylamide (PAAm) hydrogel is integrated into silicone rings. Because of the buoyance of the silicone rings, the system rises to the liquid surface to form an ALI yet can be magnetically sunk to submerged culture. However, the operation of the microboat ALI culture system requires careful manipulation of the magnetic field. Additionally, the reported PAAm hydrogel thickness is 750 μm, which is much larger than the typical upper limit of 200 μm for sufficient nutrient transport for cell growth (36). Consequently, there is a need for a facile and robust ALI culture system with tunable substrate stiffness matching that of healthy and diseased lung tissues.

Here, we design and fabricate an *in vitro* culture system by coating a thin layer of PAAm hydrogel with a thickness of 100 μm onto the classical ALI culture system, as schematically illustrated in **Fig. 1b**. The stiffness of PAAm hydrogel can be precisely controlled to match that of healthy and fibrotic airway tissues. We determine the optimum gel thickness that does not impair the transport of nutrients and biomolecules essential to cell growth. Similar to the conventional ALI system, the gel-ALI system allows HBECs to proliferate and differentiate into a pseudostratified epithelium, capturing the essential biological features *in vivo*. Using the gel-ALI system, we reveal that hydrogel substrates with pathologically relevant stiffness promote the migration of airway epithelial cells, highlighting the importance of the mechanical environment for the behavior of cells. The developed gel-ALI system represents the simplest possible *in vitro* model that enables mimicking the mechanical environment of human lung tissues, providing a platform for studying previously less explored mechanical cues on human airway biology and pulmonary diseases.

## 2. Results and Discussion

### 2.1 Design and fabrication of the gel-ALI system

We design the gel-ALI system by coating the classic ALI system with a thin layer of PAAm hydrogel. This approach represents perhaps the simplest yet unexplored method for mimicking tissue stiffness. We choose PAAm as the hydrogel coating because it has been extensively used as a substrate for studying cell mechanics (37). Moreover, it can be easily synthesized using a one-step free radical polymerization. In a typical synthesis, we fix the volume ratio between the acrylamide (AAm) monomer (40% w/v) solution and the crosslinking agent *N,N’*-methylenebisacrylamide (BIS) (2% w/v) solution at 1:1. The mixture is dissolved in DI water at a prescribed concentration. In the presence of the catalyst tetramethylethylenediamine (TEMED) and the initiator ammonium persulfate (APS), the precursor monomers polymerize to form a network, as illustrated in **Fig. 2a**. By varying the concentration of the AAm/BIS mixture, the stiffness of the PAAm gel can be tuned in a wide range from ∼80 Pa to ∼21 kPa, covering the stiffness of healthy (∼1 kPa) and IPF (∼16 kPa) lung tissues, as shown in **Fig. 2b**.

**Figure 2.**
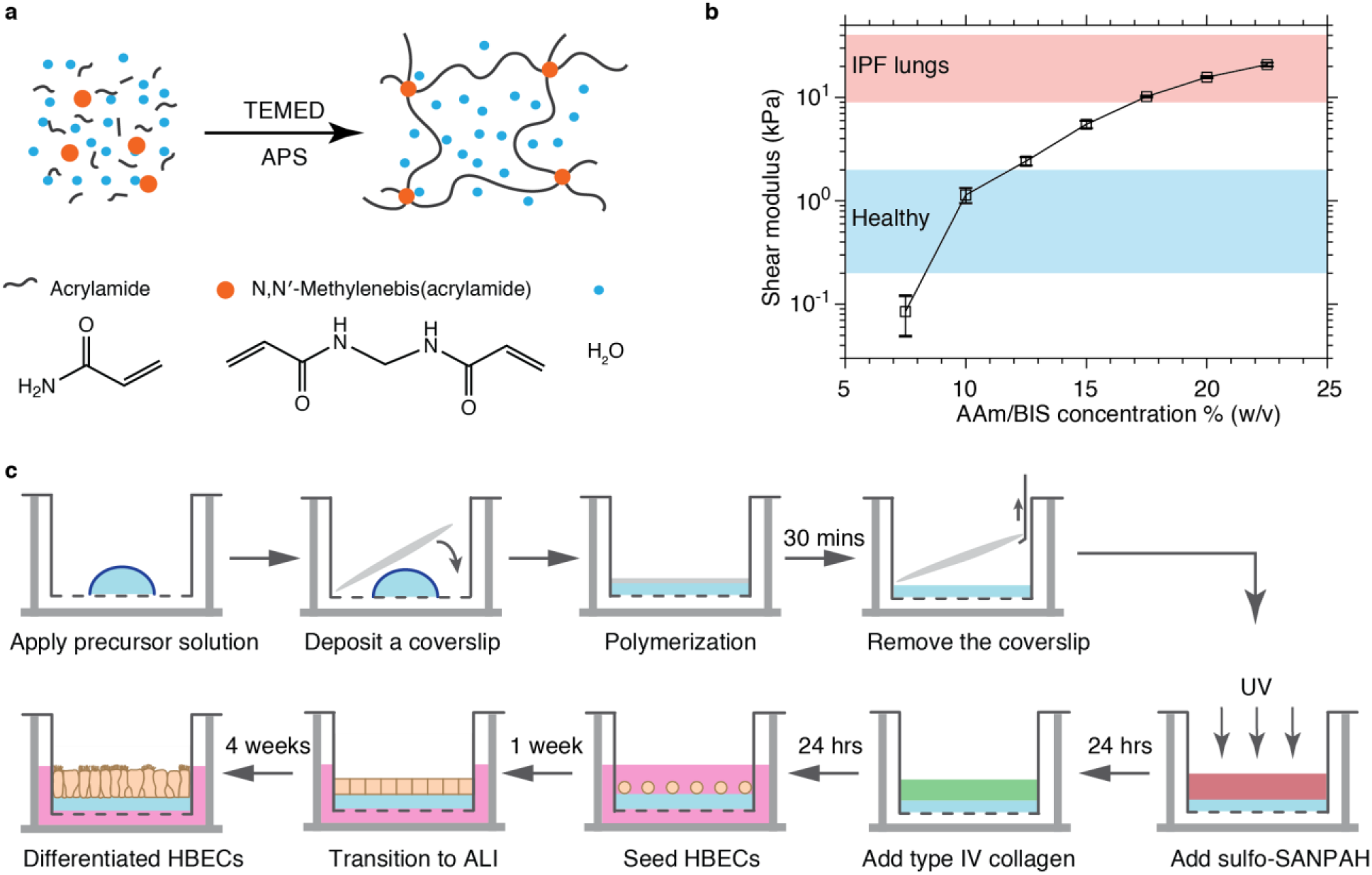
Design and fabrication of the gel-ALI culture system. **(a)** PAAm hydrogel is formed by copolymerizing AAm monomers and crosslinking agent BIS in the presence of initiator APS and catalyst TEMED. **(b)** Dependence of the shear modulus of the hydrogel on the concentration of AAm/BIS mixture measured at 20 °C at a fixed strain of 0.5% and oscillatory shear frequency of 1 rad/sec. **(c)** Flowchart for the fabrication of PAAm hydrogel-coated ALI culture system.

To fabricate the gel-ALI culture system, we apply a prescribed volume of the reaction mixture to the apical side of a Transwell^®^ insert. The solution is covered by a customized coverslip that fits the insert. Moreover, the surface of the coverslip is hydrophobic; this avoids wetting-induced spreading of the precursor hydrogel solution, such that the thickness of the hydrogel can be precisely controlled. After the solution is cured for about 30 min, we gently remove the coverslip. To quantify the gel thickness, we premix the reaction mixture with fluorescein isothiocyanate (FITC)-labeled dextran and use confocal microscopy to measure the thickness of the fluorescence region (**Materials and Methods**). This *in situ* polymerization process ensures seamless contact and strong adhesion between the soft hydrogel and the stiff plastic membrane; this is essential to prevent the delamination of the hydrogel and the leakage of cells during long-term cell culture.

### 2.2 Determine the gel thickness for efficient nutrient transport

In a gel-ALI culture, nutrients must transport through the hydrogel from the basal to the apical side to maintain cell growth. Thus, it is critical to determine the conditions for efficient mass transport through the gel. The transport is largely determined by diffusion, a process dependent on both the mesh size and the thickness of the hydrogel. The mesh size 𝜉 of a hydrogel is largely determined by the gel shear modulus 𝐺, 𝜉 ≈ (𝑘_𝐵_𝑇/𝐺)^1/3^, in which 𝑘_𝐵_ is Boltzmann constant and 𝑇 is the absolute temperature (33) (38). Because the stiffness is determined by the target lung tissues, 𝜉 is pre-determined: for the gel mimicking healthy lung tissue with 𝐺 ≈ 1𝑘𝑃𝑎, 𝜉_ℎ_ ≈16 nm; by contrast, for the gel mimicking IPF disease with 𝐺 ≈ 16 𝑘𝑃𝑎, 𝜉_𝑑_ ≈ 6 nm. Therefore, we seek to determine the gel thickness allowing for efficient nutrient transport.

We measure the time it takes for protein mimics to transport through a hydrogel layer of various thickness from 100 μm to 1000 μm. We use 70 kDa dextran as the probe molecule, which is a branched polymer with a hydrodynamic diameter of 6.4 nm (8); this value is comparable to the size of albumin, the largest protein in culture media (39). Because dextran molecules are prone to concentrate onto the surface of the plastic membrane due to unspecific interactions, we measure the fluorescence intensity of regions >25 µm above the membrane to avoid the effects of concentrated dextran on measurements, as illustrated in **Fig. 3a**.

**Figure 3.**
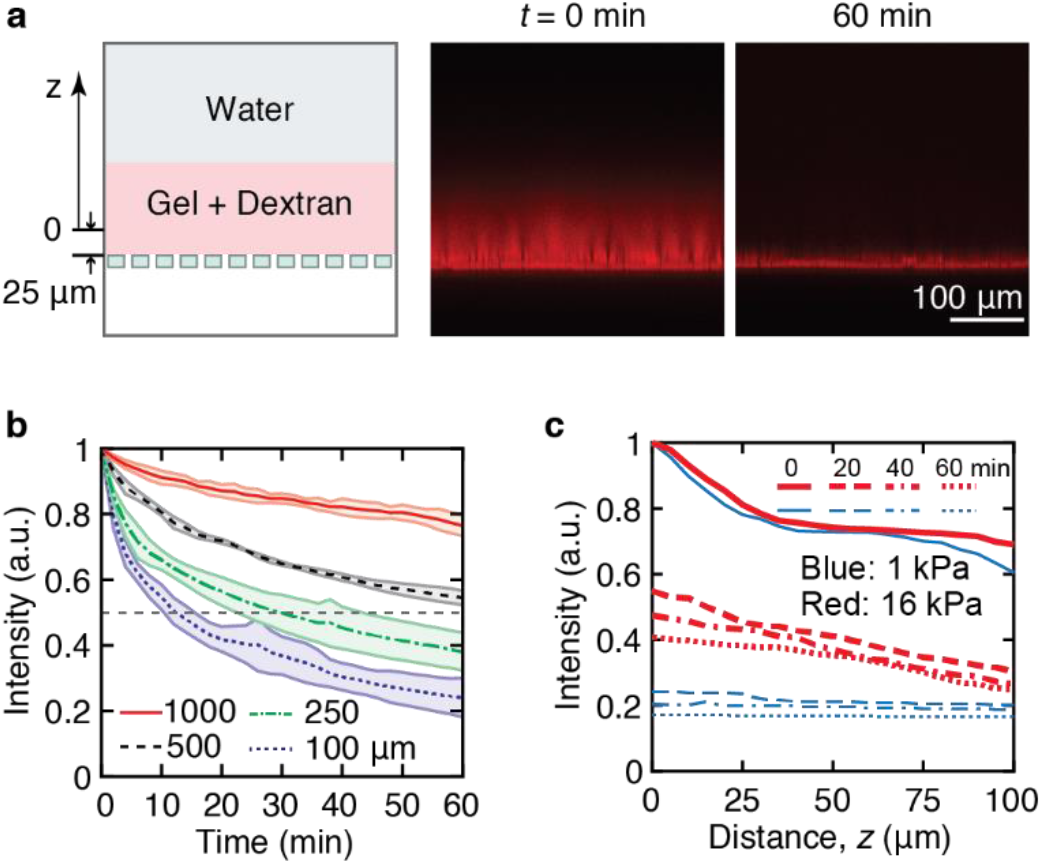
Transport properties of the gel-ALI system. **(a)** A schematic of the experimental setup (left) for measuring the diffusion of FITC-dextran molecules out of the gel. Dextran is premixed with the precursor gel solution during fabrication. *z*, the distance away from the reference point, which is chosen to be 25 μm above the porous membrane to avoid its effects on fluorescence intensity. Representative fluorescence images for 1 kPa gel of 100 μm in thickness at 0 min (middle) and 60 min (right). **(b)** Dependence of the average fluorescence intensity of the gel layer of various thicknesses on waiting time: 1000 μm (red solid line), 500 μm (black dashed line), 250 μm (green dash-dotted line), and 100 μm (purple dotted line). The measurements are performed for the region at distance >25 μm above the porous membrane. The shadowed zone represents with the width of standard deviation; *n=3*. **(c)** Fluorescence intensity profiles of the gel layer at various waiting times for gels of 1 kPa (red) and 16 kPa (blue).

To describe the transport properties of the gel, we introduce the half-decay time 𝜏, at which the fluorescence intensity of the gel decreases to 50% of the initial value. For 1 kPa gel, as the thickness increases from 100 µm to 250 µm, 𝜏 increases by nearly three times from 10 to 30 min, as shown by purple and green curves in **Fig. 3b**. Further increasing the gel thickness to 500 µm dramatically prolongs the value 𝜏 to nearly 60 min (black dashed line, **Fig. 3b**); this time is much longer than the typical nutrient supply rate required for optimal cell culture (40). As the gel thickness becomes 1000 µm, the intensity decreases to 75% of its initial value after one hour (red dotted line, **Fig. 3b**). These results confirm the understanding that thinner gels permit more efficient mass transport. Moreover, even for the 16 kPa gel, 100 µm thickness allows for relatively efficient mass transport with τ≈20 min, as evidenced by the fluorescence intensity profiles across the gel layer at different time scales (thick red dashed line in **Fig. 3c**). The determined optimum gel thickness, ∼100 µm, for efficient mass transport is much smaller than 750 µm, the thickness used for the same PAAm hydrogel in a recently developed magnetic microboat ALI culture (35). Yet, our results are consistent with the existing understanding that in native tissues mammalian cells are typically located within 100-200 µm of blood vessels (41) and that in engineered tissues only cells up to a distance of 200 µm away from the media have access to sufficient nutrients (36).

To complement the studies on transport properties of gels based on the diffusion of protein mimics, we quantify the effects of gel coating on the electrical resistance of the porous plastic membrane. The electrical resistance reflects the ionic conductivity across the gel coated membrane; therefore, it provides an indirect assessment of the transport properties of the gel. We start with determining the electrical resistance of the semipermeable membrane without gel coating, which has a value of 159±10 Ω cm^2^ that is consistent with literature data (5). When the gel is relatively thin of 100 μm, the increase in electric resistance is very small, ∼5% regardless of gel stiffness (**Fig. S1**). Collectively, our results show that 100 µm gel coating is thin enough for efficient nutrient transport.

### 2.3 Gel-ALI system allows HBECs to differentiate into pseudostratified epithelia

Despite the efficient transport of protein mimics, the ultimate test of the gel-ALI system is to examine whether HBECs can grow and differentiate into a pseudostratified bronchial epithelium. To this end, we activate the surface of the hydrogel using sulfosuccinimidyl 6-(4’-azido-2’-nitrophenylamino)hexanoate (sulfo-SANPAH), such that the hydrogel surface can be conjugated with type IV collagen, a standard ECM protein for culturing primary HBECs (**Fig. 2b**) (2). Similar to the classical ALI culture, primary HBECs are plated and cultured by adding medium to both the apical and basal sides for about one week to reach >90% confluence. Afterward, HBECs are transitioned to ALI by removing the apical medium and maintaining medium on the basal side only. The cells are cultured for 4 weeks after which HBECs are expected to fully differentiate to a pseudostratified epithelium (2).

We perform immunostaining to identify the population of cells cultured on substrates of various stiffness (**Materials and Methods**). As expected, for the control group without gel coating, HBECs fully differentiate into a pseudostratified epithelium consisting of ciliated cells and mucus-secreting goblet cells, as shown by the fluorescence images for each cell type in **Fig. 4a**. Quantitatively, for the control group the fraction of ciliated cells is 0.29±0.17, whereas the fraction of goblet cells is 0.13±0.08. These values are consistent with literature data that for *in vitro* HBEC culture the fractions of ciliated and goblet cells, respectively, are about 0.30 and 0.10-0.15 (42, 43).

**Figure 4.**
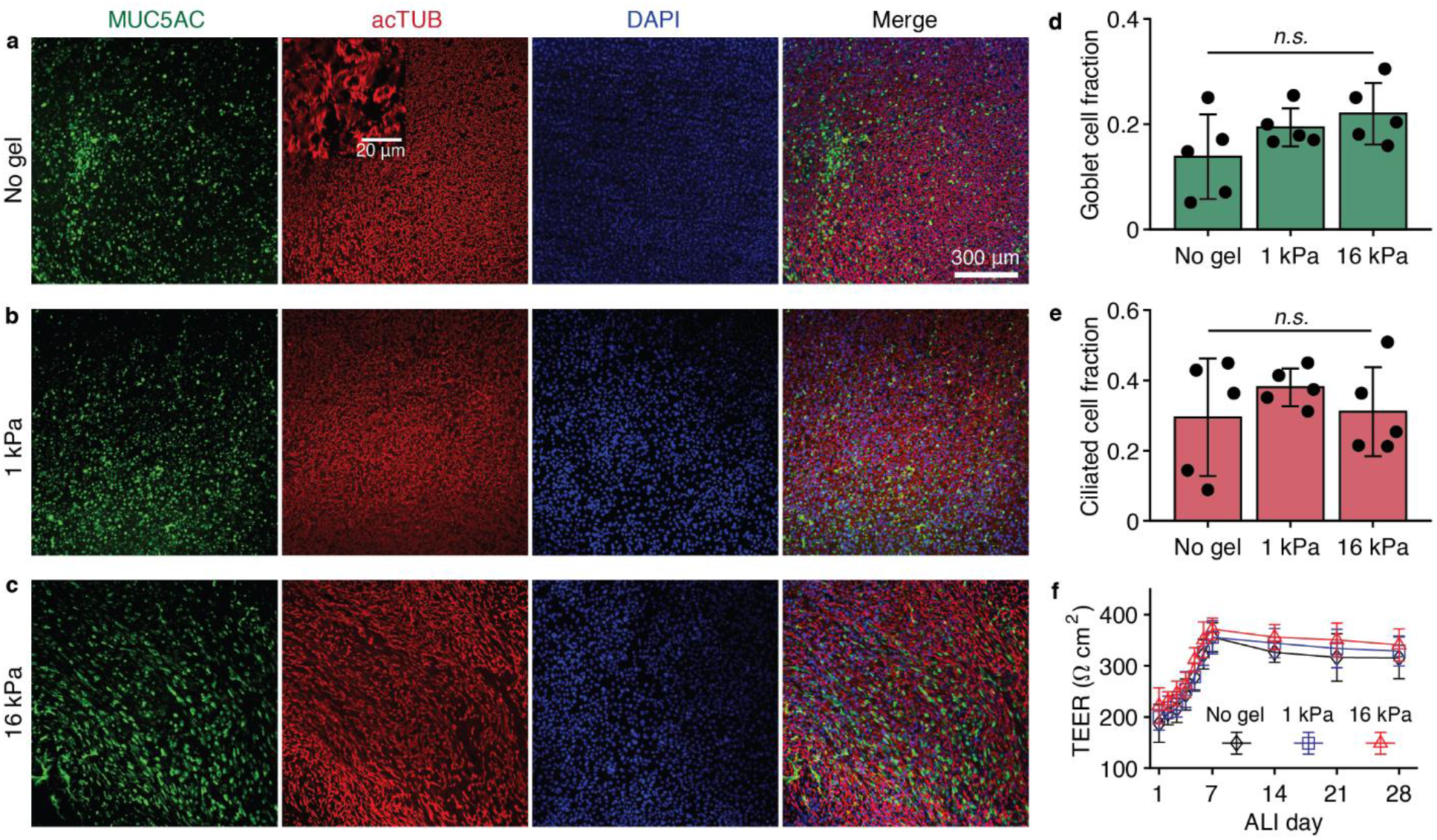
The gel-ALI system allows for HBECs differentiating into pseudostratified epithelia. **(a-c)** Immunofluorescence staining of cells after ALI culture for 4 weeks: **(a)** conventional ALI culture without gel coating, **(b)** ALI culture on 1 kPa gel, and **(c)** ALI culture on 16 kPa gel. Green: MUC5AC/goblet cells; red: AcTUB/ciliated cells; blue: DAPI/nuclei. Inset in **(a)**: Zoom-in view of cilia. The fraction of **(d)** ciliated cells and **(e)** goblet cells on substrates of various stiffnesses. **(f)** Transepithelial electrical resistance (TEER) of HBEC cultures at different ALI days. *n.s.*, not significant. Values are mean ± SD, *n* = 5 different donors. One-way ANOVA is used for statistical analysis.

Similarly, HBECs differentiate into pseudostratified epithelia on gels of 1 kPa (**Fig. 4b**) and 16 kPa (**Fig. 4c**). Interestingly, regardless of gel coating, the fraction of ciliated cells is in the range of 0.3-0.4 (**Fig 4d**). Moreover, the fraction of goblet cells is around 0.2 and there is no significant difference among substrates of various stiffnesses (**Fig. 4e**). The stained ciliated and goblet cells together account for about 50% of the whole cell population; the rest unstained ones are other cell types such as basal cells and club cells (44, 45).

In parallel, we quantify the transepithelial electrical resistance (TEER) of HBEC cultures, a well-established method used to monitor HBECs during their various stages of growth and differentiation (46). We subtract the contribution from the gel coated semipermeable membrane to obtain the TEER values for HBECs. Up to ALI Day 7 the HBEC TEER value increases nearly linearly from ∼200 to 350 Ω cm^2^, reflecting the proliferation of HBECs (**Fig. 4f**). After ALI Day 7, the TEER value slightly decreases and becomes stable beyond ALI Day 21, reflecting the complete differentiation of HBECs. Moreover, the plateau TEER value 329±33 Ω cm^2^ is consistent with previously reported ones (5) for well differentiated HBECs with tight cell junctions (**Fig. 4f**). Taken together, our results show that gel-ALI allows HBECs to differentiate into a pseudostratified epithelium.

### 2.4 Pathological substrate stiffness promotes cell migration

We exploit the gel-ALI system to study the migration of HBECs, a process critical to human airway remodeling (47). Three groups are explored: (i) control with the conventional plastic substrate, (ii) hydrogel with stiffness of 1 kPa matching that of healthy lung tissues, and (iii) hydrogel with stiffness of 16 kPa matching that of fibrotic lung tissues. For each group, we use cells from five donors to ensure statistical significance. The difference in refractive indices between the cell junction and the cell body allows us to use phase-contrast microscopy to measure the migration of cells (**Materials and Methods**). And we measure cell migration on different ALI days up to 3 weeks, a typical period for basal cell differentiation, as shown by representative images in **Fig. 5a** and **Movies S1-S3**. Using particle image velocimetry (PIV) (48), we compute the velocity map of cells, as shown by the heat map in **Fig. 5b** and **Movies S1-S3**.

**Figure 5.**
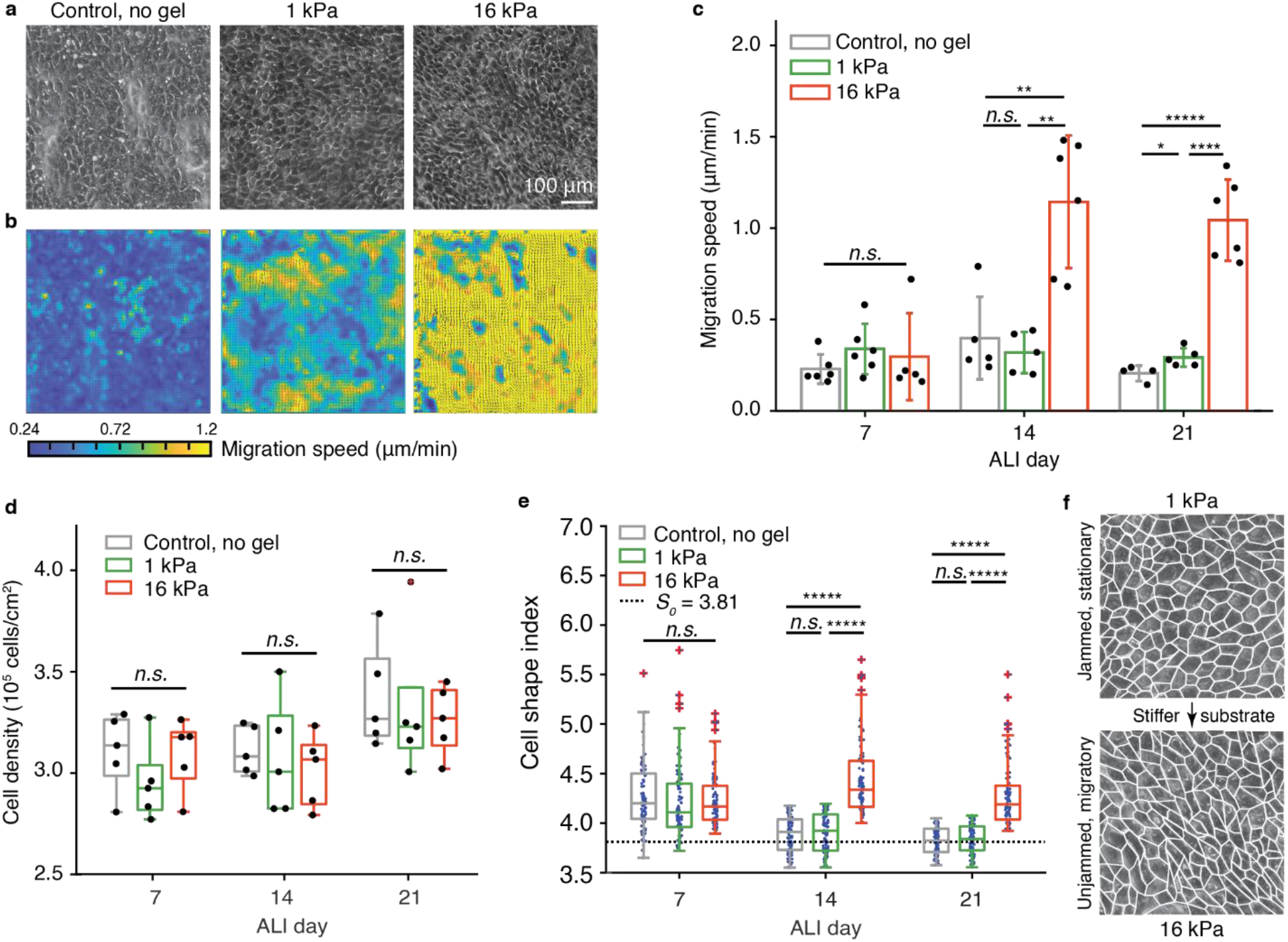
A stiffer substrate promotes migration of HBECs. **(a)** Example phase-contrast images of HBECs cultured on substrates of different stiffness on ALI day 21. **(b)** Speed maps for the migration of HBECs on substrates of different stiffness on ALI day 21. **(c)** Average cell migration speed, **(d)** density, and **(e)** cell shape index *S* of HBECs under different substrate stiffness on ALI days 7, 14, and 21. *n.s.*: not significant; * *p* < 0.05, ** *p* < 0.01, *** *p* < 0.001, **** *p* < 0.0001, ***** *p* < 0.00001. For **(c)**, values are mean ± SD, *n* = 4-6 donors; for **(d)**, *n* = 5 donors; for **(e)**, *n* =100 cells. One-way ANOVA is used for statistical analysis. **(f)** Substrates with pathological stiffness promote migration of cells by fluidizing jammed epithelia.

The effects of gel stiffness on cell migration become more pronounced on longer ALI days. For example, on ALI day 7, there is no significant difference in cell migration speed regardless of substrate stiffness, as shown by the histograms on the left of **Fig. 5c**. However, on ALI day 14, despite no significant difference between control and 1 kPa gel in cell migration speed (∼0.30 μm/min), the speed for HBECs on 16 kPa gel increases by nearly 4 times to 1.14 μm/min. Moreover, a similar trend is observed on ALI day 21, although HBECs on 1 kPa gel migrate slightly faster than the control, as shown by the histograms on the right of **Fig. 5c**. Yet, across the 21 days, there is no significant difference in cell migration speed among ALI day 7, 14, and 21 for the control and 1 kPa gel, and only cells on 16 kPa gel exhibit markedly high migration speed starting ALI day 14.

Because the migration of cells in a confluent monolayer is known to be slower at a higher cell density (49, 50), we quantify the cell density under various substrate stiffness. For each cell culture, we count the number of cells per unit area and perform the same measurements for cells from five donors to ensure sufficient statistics. We find no significant difference in cell density over the 21-day period, as shown in **Fig. 5d**. Moreover, the average cell density is (3.2 ± 0.2) × 10^5^/𝑐𝑚^2^, consistent with the previously reported value (42). These results preclude that the observed difference in cell migration speed is attributed to cell density.

To explain the difference in cell migration speed, we exploit a recently developed theoretical framework that correlates the migration of a confluent epithelial monolayer with cell shape (51, 52). The theory predicts that the cell shape index, 𝑆 = 𝑃/√𝐴, where 𝑃 is the cell perimeter and 𝐴 is the cell area, can be an indicator for cell migration. Below a threshold value with 𝑆 < 𝑆_0_ = 3.81, cells are more round-like and are jammed, stationary; by contrast, for 𝑆 > 𝑆_0_, cells are more elongated and are unjammed, migratory. Consistent with this understanding, on the same ALI day there is no significant difference in 𝑆 between cells cultured on the control and the 1 kPa gel on ALI day 7 and day 14; by contrast, for the pathologically relevant hydrogel of 16 kPa the value of *S* is significantly higher at 4.2 (**Fig. 5e**) on ALI day 14. Moreover, on ALI day 21 when the HBECs on the 1 kPa gel are nearly stationary, the cell shape index becomes 3.82, comparable to the threshold value 𝑆_0_, while the cell shape index of HBECs on 16 kPa is still significantly higher at 4.1 (**Fig. 5e**). These results indicate that hydrogel substrate of pathological stiffness promotes the migration of HBECs at the air-liquid interface, highlighting the potential of the gel-ALI system for studying the effects of mechanical cues on fibrotic lung diseases.

## 3. Conclusion

We have developed a gel-ALI culture system by coating a thin layer of polyacrylamide hydrogel onto the porous plastic membrane of the Transwell^®^ insert of the classic ALI culture system. The hydrogel stiffness can be tuned in a wide range to precisely match that of the healthy and fibrotic lung tissues. To allow efficient nutrient transport, the hydrogel must be relatively thin at ∼100 μm, significantly lower than the 750 μm gel thickness in a recently reported magnetic hydrogel boat-based ALI system (35). We show that the gel-ALI system allows primary HBECs to fully differentiate into a pseudostratified epithelium, capturing essential biological features of human airway tissues *in vivo*.

Using the gel-ALI system, we studied the effects of substrate stiffness on the migration of HBECs, a process critical to human airway remodeling. Moreover, we discover that hydrogel substrates of pathologically relevant stiffness promote the migration of HBECs at air-liquid interface; this represents a previously unrecognized phenomenon but can be rationalized in the context of a theoretical framework that associates jamming/unjamming transition of a confluent epithelial layer to the cell shape index.

Compared to the recently developed microfluidics-based (53) or magnetic hydrogel boat-based (35) ALI systems that rely on sophisticated microfabrication that often is not accessible to biological laboratories, our gel-ALI system is built on the classic, commercially available ALI system that has been widely used in biological laboratories. In addition, the polyacrylamide hydrogel chemistry is well established, and the fabrication process is facile and does not require any sophisticated instrument. Moreover, integrating another polymer network such as alginate into the polyacrylamide hydrogel can result in a double-network hydrogel (54, 55); this would enable controlling not only hydrogel stiffness but also relaxation time to mimic the dynamic nature of biological tissues, which are being increasingly recognized to be critical to fundamental cellular processes (56). Finally, the gel-ALI system is not restricted to culture HBECs; instead, it should be general and applicable to small airway cells, which are more relevant to fibrotic lung diseases (57–59). In addition, the gel-ALI platform may be applied to culture intestinal epithelial cells (60) to study the effects of fibrosis on intestinal epithelial cells in the context of inflammatory bowel diseases (61, 62). Thus, we believe that the developed gel-ALI system represents perhaps the simplest yet versatile platform for studying the roles of pathologically relevant mechanical cues in human airway biology and diseases.

## Materials and Methods

### Fabrication and characterization of gel-ALI system

To synthesize polyacrylamide (PAAm) hydrogel with a prescribed stiffness, we prepare a reaction mixture mixing acrylamide solution (AAm, MilliporeSigma, Cat. No. A4058, 40% w/v), *N,N’*-methylenebisacrylamide solution (BIS, MilliporeSigma, Cat. No. M1533, 2% w/v), and deionized (DI) water (Millipore), following the recipe listed in **Table S1**. To initiate polymerization, a final concentration of 0.12% w/v ammonium persulfate (APS, MilliporeSigma, Cat. No. 248614) is added as the catalyst, and 1.2% v/v N,N,N’,N’-tetramethylethylenediamine (TEMED, MilliporeSigma, Cat. No. T9281) as an initiator is added to the mixture.

To coat a Transwell^®^ insert (Corning, Cat. No. 3401) with a PAAm hydrogel layer of 100 μm thickness, we quickly apply 30 μL of the reaction mixture to the center of the Transwell^®^ insert. In parallel, we prepare a circular coverslip of 11 mm in diameter by punching three layers of Scotch^®^ tape and use the coverslip to spread the reaction mixture. Because the coverslip is hydrophobic and has a diameter slightly smaller than that of the Transwell^®^ insert which is 12 mm, this process flattens the gel surface, and importantly, prevents the reaction mixture from oxygen, which is known to impair free radical polymerization. After 30 min, the coverslip is removed by a hooked syringe, leaving a thin layer of hydrogel with the prescribed thickness. Afterward, the gel is rinsed using sterilized DI water. We seal 12-well plates containing gel-coated Transwell^®^ inserts by parafilm and store them at 4 °C for future usage.

To quantify the gel thickness, we add 50 μg/mL fluorescein isothiocyanate (FITC)-labeled dextran with a molecular weight of 70 kDa (Invitrogen, Cat. No. D1823) to the reaction mixture. The gel thickness is quantified by measuring the thickness of the fluorescent region.

### Rheological characterization of polyacrylamide hydrogel

We prepare a gel disk for rheological measurements. To do so, we add 800 μL reaction mixture to a well in a 12-well culture plate and cover the solution using a circular glass coverslip of 21 mm in diameter. After 30 min, the coverslip is removed, leaving a gel layer of ∼2 mm in thickness. Form the gel layer, we cut a disk of 8 mm in diameter, load it to a stress-controlled rheometer (Anton Paar, MCR 302) with a plate-plate geometry of 8 mm in diameter, and cover the gel peripheral with mineral oil to avoid evaporation. We quantify the dynamic mechanical properties using oscillatory shear measurements at a fixed strain of 0.5 % but various frequency of 0.1 rad/sec to 100 rad/sec. Within the frequency range explored, the shear storage modulus is nearly independent of the shear frequency. Therefore, we take the shear storage modulus at the lowest shear frequency, 0.1 rad/sec, as the equilibrium shear modulus *G* of the gel.

### Measurement of transport properties gel-ALI system

We label the gel by premixing the reaction mixture with 70 kDa FITC-labeled dextran at a concentration of 50 μg/mL. The gel-ALI system is fabricated following the same protocol described above. We add DI water to the apical chamber of the Transwell^®^ insert and monitor the fluorescence intensity using a fluorescence confocal microscope (Leica, SP8). The fluorescence intensity is quantified and normalized to the value at the initial time point.

### Cell culture

We coat the gel-ALI system with type IV collage for HBEC culture. We activate the gel surface by sulfosuccinimidyl 6-(4’-azido-2’-nitrophenylamino) (sulfo-SANPAH, Thermo Scientific, Cat. No. 22589), which allows covalent conjugation of type IV collagen to the gel surface. Specifically, to each gel-coated Transwell^®^ insert we add 100 μL of 2 mg/mL sulfo-SANPAH (MilliporeSigma, Cat. No. D2650) solution in DI water, expose the solution to UV light (37 mW, 365 nm) for 5 min to activate the gel surface, and then rinse the gel using DI water. For each well with activated gel surface, we add 150 μL of 0.005 mg/mL type IV collagen (MilliporeSigma, Cat. No. C7521) solution in DI water and store the gel-ALI system at 4 °C overnight to ensure adequate collagen coating. Then, the gel is sterilized by UV light for 15 min and ready for cell culture.

Primary HBECs are obtained from Marsico Lung Institute Tissue Procurement and Cell Culture Core at the University of North Carolina at Chapel Hill. For all experiments, we use passage 2 cells, beyond which the HBECs may lose their stemness (2). We seed HEBCs at the density of 4.2×10^4^ cells/insert and add expansion medium (PneumaCult™-Ex Plus Medium, STEMCELL Technologies, Cat. No. 05040) to both the basal and apical chambers. After 5-7 days, HBECs reach over 90% confluence. We transition the cultures to ALI by removing the apical medium and replacing the expansion medium with ALI culture media (PneumaCult™-ALI Medium, STEMCELL Technologies, Cat. No. 05001). After 2 weeks of ALI culture, mucus starts to accumulate and is washed three times per week. To wash off mucus, to each insert we add 500 μL of Dulbecco’s phosphate-buffered saline (DPBS, Gibco 14-200-075), incubate the cell culture for 15 min, and aspirate the DPBS in the apical chamber. The washing process is repeated three times.

### Immunofluorescence microscopy

We use immunofluorescence staining to label the cells in a well-differentiated HBEC culture. To prepare for the staining, for each Transwell^®^ we wash the culture three times using DPBS; for each washing, 500 μL of DPBS is added to the apical chamber and the culture is incubated at 37 ℃ for 20 min. After washing, we add 1 mL of 4% w/v paraformaldehyde (Thermo Scientific, Cat. No. AA47392-9M) to the apical side of the Transwell^®^ and incubate the culture at room temperature (RT) for 15 min to fix the cells. Then, we add 1 mL of 0.2% v/v Triton X-100 solution (Thermo Scientific, Cat. No. AAA16046AP) and incubate the culture at RT for 30 min to permeabilize the fixed cells. We add 1 mL of 3% (v/v) donkey serum (Abcam, Cat. No. AB7475) in DPBS solution and incubate the culture at RT for 1 hour; this process blocks non-specific binding of antibodies to cells.

For the above prepared cell culture, we use mouse anti-mucin 5AC (MUC5AC) antibody (ThermoFisher, Cat. No. MA5-12178) to label mucin granules in goblet cells and rabbit anti-acetylated tubulin (AcTUB) antibody (Invitrogen, Cat. No. MA5-33079) to label tubulin in the cilia of ciliated cells. Specifically, 200 μL of primary antibody solution containing 10 μg/mL anti-MUC5AC antibody and 40 μg/mL of anti-AcTUB antibody is added to each well. The culture is maintained at 4 ℃ overnight to ensure the binding of the primary antibody to antigens. Afterward, we rinse the cell culture with DPBS three times and add to each well 200 μL of secondary antibody solution, which contains 10 μg/mL donkey anti-rabbit secondary antibody conjugated with Alexa Fluor 647 (Abcam, Cat. No. ab150075) and 10 μg/mL donkey anti-mouse secondary antibody conjugated with Alexa Fluor 488 (Abcam, Cat. No. ab150105). We incubate the culture at RT for 1 hour to allow the secondary antibodies to bind to the primary antibodies, and rinse the culture with DPBS three times afterward. After staining using secondary antibody, to each well we add 200 μL of 1 μg/mL 4’,6-diamidino-2-phenylindole (DAPI, Invitrogen, Cat. No. D3571) solution in DI water, incubate the culture at RT for 5 min, and then wash the culture with DPBS three times. This process is to stain the cell nuclei.

To prepare the culture for imaging, we excise the Transwell^®^ insert membrane, mount it on a glass cover slide, add about 1 mL FluorSave oil (MilliporeSigma, Cat. No. 345789) to completely cover the membrane, place a coverslip, and seal the edges of the coverslip using wax. We use multi-channel fluorescence confocal microscopy (Leica SP8) to image the cell culture. During imaging, we use sequential scan between frames to avoid the overlap of fluorescence from different channels. For each code of cell culture, we randomly select five regions of interest (ROIs) from five wells with one ROI from each well. We use five codes of cells from different healthy donors to ensure statistical significance. For each culture, we use a 10× dry objective to image a ROI with the dimension of 1162.5×1162.5 μm^2^; this allows us to image about 4000 cells/culture and a total number of ∼10^5^ cells for each condition to avoid biased measurement of the cell population. To image the cilia of ciliated cells for the control sample, we use a 63× oil objective (numerical aperture 1.4) to obtain a high-resolution image of individual cilia (inset, **Fig. 4a**). The fraction of each cell type is calculated by dividing the cell area by the total area of the image.

### Transepithelial electric resistance (TEER) measurement

We measure TEER using an EVOM2 epithelial voltohmmeter (World Precision Instruments, Sarasota, FL). Before each measurement, we sterilize the electrodes with 70% ethanol, rinse them using sterilized DI water, and dry them. For each culture, we add 100 μL of DPBS to the apical side of the Transwell^®^ to form an electrical circuit with the liquid in the basal chamber. We first measure the electric resistance of a blank Transwell^®^ with or without gel coating. To measure the electric resistance across the HBEC culture, we gently place the electrodes into the medium on both the apical and basal sides of the cell culture without disrupting the cell surface. To obtain TEER for HBECs, the measured resistance is subtracted by the value of a blank Transwell^®^ with or without gel and multiplied by the area of Transwell^®^. The measurements are performed on ALI days 1, 2, 3, 4, 5, 6, 7, 14, 21, and 28 to quantify the dependence of TEER values on culture time.

### Quantification of HBEC migration and cell density

To monitor the migration of HBECs, we use a confocal microscope equipped with a 10x dry objective to acquire phase contrast images. The phase contrast mode allows for detecting the differences in the refractive index of cell body and junction, enabling label-free imaging of HBECs at their natural state. Moreover, the confocal microscope is equipped with an environmental chamber with temperature, CO_2_, and humidity control to allow for long time imaging. For each condition, we image HBECs at 1 frame/min for 60 min on ALI days 7, 14, and 21. We use particle imaging velocimetry (PIV) to quantify the velocity fields associated with collective cell migration (48). Interrogation windows of 64×64 pixels with 50% overlap are used; this window size allows for a spatial resolution of ∼16 pixels (9 μm).

To measure cell density, for a randomly selected ROI, we count the number of cells manually. For cells on the four borders of the ROI, only cells on the upper and left borders are included. The cell density is calculated as the total number of cells divided by the area of ROI. For all the measurements, we use cells from 4-6 healthy donors to ensure statistical significance.

### Measurement of the cell shape index

The cell shape index is defined as 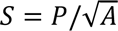, where 𝑃 is the cell perimeter and 𝐴 is the cell area. We use ImageJ to trace the cell contour to measure the cell perimeter and area to calculate the cell shape index. For each condition, we randomly choose 100 cells from 5 wells representing 5 healthy donors to ensure statistical significance.

### Statistical analysis

Statistical analysis is performed using the one-way analysis of variance (ANOVA) method.

## Acknowledgments

L.H.C. acknowledges the support from NSF (CAREER DMR-1944625). We thank Dr. Kenichi Okuda at UNC Chapel Hill for providing cells. We thank Dr. Kenichi Okuda, Dr. Gang Chen, Dr. Camille Ehre, and Dr. Lin Sun at UNC Chapel Hill for helpful discussions. We thank Dr. Alison Criss from UVA for the assistance with the TEER measurements.

## Author contributions

L.H.C., Z.J.H, and C.C. designed the research. Z.J.H. developed the protocol to fabricate the gel-ALI system. Z.J.H. and C.C performed rheological measurements of gel stiffness and analysis. C.C measured the gel transport properties. Z.J.H and R.D cultured HBECs on gel-ALI and performed immunostaining. C.C cultured HBECs on gel-ALI and measured the cell migration speed, the cell density, and the cell shape index. L.H.C., Z.J.H., and C.C. wrote the paper. All authors reviewed and commented on the paper. L.H.C. conceived and supervised the study.

## Supplementary Material

<u>gelALI_SM.pdf</u>

https://figshare.com/s/a2e2951a3cf71fef5c8d

## Supplementary Movies

HBEC migration on substrates of different stiffness.

https://figshare.com/s/f39a7c6096005d4553e2

