## Supplemental Materials for "A gel-coated air-liquid-interface culture system with tunable substrate stiffness matching healthy and diseased lung tissues"

**This PDF file includes**

Figure S1
Table S1

**Supplementary Text**

To complement the studies on transport properties of gels based on the diffusion of protein mimics, we quantify the effects of gel coating on the TEER of the porous plastic membrane. The TEER reflects the ionic conductivity across the gel coated membrane; therefore, it provides an indirect assessment to the transport properties of the gel. We start with determining the TEER of the semipermeable membrane without gel coating, which has a value of 159±10 Ω that is consistent with literature data (1). Introducing gel coating increases the electric resistance, and the increase is nearly linearly proportional to the thickness of gel coating; this is consistent with the understanding that the ionic conductance decreases linearly with the increase of gel thickness (left solid line, **Fig. S1**). Yet for thick gels greater than 500 μm, the electric resistance saturates around 170 Ω, about 20% increase compared to the membrane without gel coating (right solid line, **Fig. S1**). Moreover, for gels of the same thickness, the stiffer gel has slightly higher electric resistance (**Fig. S1**); this is because the stiffer gel has a smaller pore size and thus lower ionic conductivity. Nevertheless, when the gel is relatively thin of 100 μm, the increase in electric resistance is very small of ~5% regardless of gel stiffness (**Fig. S1**).

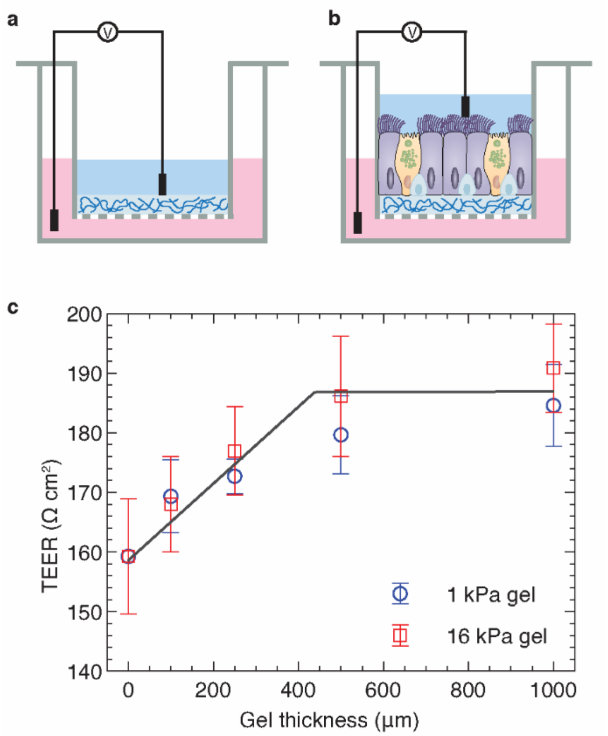


**Figure S1. Transepithelial electric resistance (TEER) of gel-ALI systems**. (a, b) Schematics for measuring TEER of empty Transwell® and human bronchial epithelial cell cultures in gel-ALI systems. (c) Dependence of TEER of the gel-ALI system without cells on gel thickness and stiffness. Values are mean ± SD, $n$ = 5.

**Table S1. Stiffness of polyacrylamide gel correspondent to different added volumes of AAm, BIS, and DI water.**

| AAm solution 40% w/v (μl) | BIS solution 2% w/v (μl) | DI water (μl) | TEMED (μl) | APS solution 10% w/v (μl) | Shear storage modulus (Pa) |
|---|---|---|---|---|---|
| 75 | 75 | 826 | 12 | 12 | 85 ± 36 |
| 100 | 100 | 776 | 12 | 12 | 1140 ± 190 |
| 125 | 125 | 726 | 12 | 12 | 2421 ± 227 |
| 150 | 150 | 676 | 12 | 12 | 5501 ± 459 |
| 175 | 175 | 626 | 12 | 12 | 10204 ± 210 |
| 200 | 200 | 576 | 12 | 12 | 15705 ± 339 |
| 225 | 225 | 526 | 12 | 12 | 20842 ± 867 |
